# Genomic prediction in insects: a case study on wing morphology traits in the jewel wasp *Nasonia vitripennis*

**DOI:** 10.1101/2024.10.09.617398

**Authors:** Shuwen Xia, Gabriella Bukovinszkine Kiss, Hendrik-Jan Megens, Martien A.M. Groenen, Bas J. Zwaan, Piter Bijma, Bart A. Pannebakker

## Abstract

**Background:** Biological control is a sustainable strategy to combat agricultural pests. Yet, due to legislation, importing non-native biocontrol agents is increasingly restricted. Thus, selective breeding of biocontrol traits of native species is suggested to enhance performance of existing biocontrol agents. Genomic prediction is a new alternative to exploit genetic variation for improving biocontrol efficacy. This study aims to establish proof-of-principle for genomic prediction in insect biocontrol agents, using wing morphology traits in the parasitoid *Nasonia vitripennis* Walker (Pteromalidae) as a model.

**Methods:** We performed genomic prediction using a Genomic Best Linear Unbiased Prediction (GBLUP) model, using a total of 1,230 individuals with 8,639 SNPs generated by genotyping-by-sequencing (GBS). We used individuals from two generations from the outbred HVRx population, 717 individuals from generation G169 and 513 from generation G172. To assess genomic prediction accuracy, we used across-generation validation: forward validation for G172 from G169, backward in time validation for G169 from G172, and also 5-fold cross-validation, randomly using one fifth of the population as validiation and the others as training groups.

**Results:** For size-related traits, including tibia length, wing length, width, and second moment wing area, the accuracy of genomic prediction was close to zero in both across-generation validations, but much higher in 5-fold cross-validation (ranging 0.54-0.68). For the shape-related trait wing aspect ratio, a high accuracy was found for all three validation strategies, with 0.47 for across-generation forward validation, 0.65 for across-generation backward validation, and 0.54 for 5-fold cross-validation.

**Conclusion:** Promising accuracies were observed for all traits in 5-fold cross-validation, but not in the across-generation validations. Overall, applying genomic selection in insect biocontrol agents with a relative small effective population size seems promising. However, factors such as the biology of insects, the techniques of phenotyping, and costs of large-scale genotyping still challenge the application of genomic selection to biocontrol agents.

## Introduction

Biological control is a strategy to combat agricultural pests using natural enemies of pest species, such as predators and parasitoid wasps, as biocontrol agents (DeBach, 1958; Van Den Bosch, 1971). Over the past decades, effective biocontrol strategies have been established, using both native and non-native natural enemy species (van Lenteren et al., 2011). However, the Nagoya Protocol on Access and Benefits Sharing (https://www.cbd.int/abs) has considerably limited the import of non-native biocontrol agents (Cock et al., 2010; van Lenteren et al., 2011; Mason et al., 2018, 2023). Thus, optimizing the use of existing and native biocontrol agents became necessary. To enhance the efficacy of existing biocontrol agents, the use of genetic tools has been discussed over the last 50 years (DeBach, 1958; Hoy, 1986; Bielza et al., 2020; Leung et al., 2020). However, the potential for genetic improvement of biocontrol populations has largely been unexplored (Wajnberg, 2004; Lommen et al., 2017). Selective breeding, which uses the presence of standing genetic variation to select for traits of interest in biocontrol populations, may offer an opportunity to improve the performance of natural enemies used in biocontrol.

Genomic selection is a promising selective breeding approach that uses information from genome-wide DNA-markers to efficiently select for complex traits (Meuwissen et al., 2001). The availability of large numbers of single nucleotide polymorphisms (SNPs) enabled the use of genomic selection, and it has been extensively studied in livestock and plant breeding over the last decade (Meuwissen et al., 2013). In genomic selection, the quantitative trait loci (QTL) affecting the traits of interest are assumed to be in linkage disequilibrium (LD) with one or more SNP markers. The predictor of an individual’s genetic merit is the genomic estimated breeding value (GEBV), obtained as the sum of all SNP marker effects of the individual. GEBVs are then used to rank selection candidates based on their genotype to select parents for the next generation of breeding without phenotypic data. Genomic selection has revolutionized animal and plant breeding, and genomic prediction is increasingly used in human genetics to predict the individual risk of complex diseases (Schaeffer, 2006; Hayes et al., 2009; Heffner et al., 2009; Jannink et al., 2010; Wray et al., 2019). So far, however, the potential of genomic selection for genetic improvement of insect natural enemies has not been reported.

As a proof-of-principle of genomic selection in biocontrol agents, here we focus on its application in a parasitoid wasp. Parasitoids are among the most widely utilized insect biocontrol agents in practice (Waage & Hassell, 1982; Strand & Obrycki, 1996). The *Nasonia* genus, gregarious parasitoid wasps of blowfly pupae, has been used for over half a century as a model species in developmental and evolutionary genetics (Whiting, 1967; King, 1993; Rivero & West, 2002, 2005; Werren & Loehlin, 2009; Werren et al., 2010; Pannebakker et al., 2011). It includes four species: *N. vitripennis*, *N. longicornis*, *N. giraulti*, and *N. oneida* (Werren et al., 2010). The best studied species in the genus is *N. vitripennis*, which has its genome sequence released and annotated. The *Nasonia* genome consists of five chromosomes with a total size of ∼335 Mb and ∼446 cM (Werren et al., 2010; Lynch, 2015). The availability of a genome assembly has driven the development of a range of genomic and genetic tools which have facilitated the investigation of gene expression and regulation, and the identification of genes involved in complex genetic traits (Werren et al., 2010, 2016; Pannebakker et al., 2013; Wang et al., 2013, 2015; Ferree et al., 2015; Lynch, 2015; Dalla Benetta et al., 2019; Buellesbach et al., 2022). The availability of a genome assembly and other genetic tools makes *N. vitripennis* an interesting model organism to investigate the prospects of genomic prediction in biocontrol agents.

The main goal of this study was to seek proof-of-principle for the use of genomic prediction in biocontrol agents, using the parasitoid wasp *N. vitripennis* as a model organism. Importantly, *N. vitripennis* is a gregarious parasitoid, which can lay up to 60 eggs in a single dipteran pupa (Whiting, 1967). This can create environmental similarity between offspring developing within the same pupa (here referred to as “host”), which needs to be accounted for in the statistical model to avoid confounding environmental with genetic effects. Moreover, we previously found that much of the phenotypic variance of wing morphology and tibia length can be attributed to the host in which the wasp had developed (Xia et al., 2020). Thus, a common environment effect (i.e., host effect) should be included in the genomic prediction analysis. To ensure the statistical power of genomic prediction, a sufficiently dense and well-distributed set of markers across the whole genome is required. In many animal and plant species, flexible low-cost high-throughput genotyping arrays have been developed and commercialized for this purpose. However, because of the limited uptake of genetic investigation of biocontrol agents, a high-throughput genotyping SNP array is not available for *N. vitripennis*. As an alternative, Genotyping-by-Sequencing (GBS) is a genotyping approach to obtain genome-wide marker genotypes from sequence data, and has been applied in a variety of breeding schemes, especially in plants (Davey et al., 2011; Elshire et al., 2011; Poland et al., 2012; Poland & Rife, 2012; Crossa et al., 2013; De Donato et al., 2013; Wu et al., 2015). Here, we investigate the prospects of genomic prediction based on GBS in insect natural enemies, using tibia length and wing morphology traits in *N. vitripennis*. To measure the quality of the resulting GEBVs, we performed different validation strategies, both within and across generations. We found that accuracy of genomic prediction differed between traits and validation strategies, which could be largely explained by the biology of the species, such as the large effect of the shared host environment, and the study populations used.

## Material and methods

### Nasonia vitripennis *stock culture*

As our source population we used the HVRx outbred laboratory population, established from strains collected from a single population in The Netherlands (van de Zande et al., 2013). To preserve genetic diversity across generations, the HVRx stock is maintained according to a fixed schedule, in which approximately 120 mated females were transferred to four new mass culture vials to initiate the next generation (van de Zande et al., 2013). Per vial, 50 blow fly pupae (*Calliphora vomitoria* Linnaeus Calliphoridae) were provided as hosts for oviposition. To ensure optimal mixing of the wasps, the parasitized hosts were re-distributed over four new mass culture vials each generation before offspring emerge. Approximately 14 days were needed to complete a cycle at 25 °C at 16 hours light : 8 dark conditions.

### Morphological trait measurements

In total, 1,248 females were randomly collected from two generations of the HVRx population; 720 individuals from generation 169 (G169), and 528 individuals from generation 172 (G172). We recorded the host identity for individuals from G172, but not for individuals from G169. The right forewing and right hind- tibia were detached and mounted in the mounting medium Euparal (Waldeck GmbH & Co. KG, Division Chroma, Münster, Germany) under coverslips on microscope slides. Slides were photographed on a Zeiss Imager.A1 microscope (Carl Zeiss AG, Göttingen, Germany) at 2.5x magnification. Data for wing morphology were obtained by positioning landmarks on each digitized wing, using tpsDig software (Rohlf, 2013) following our previous study (Xia et al., 2020). Coordinates of the landmarks were used to calculate the following wing traits: wing length, wing width, the second moment area, and the wing aspect ratio (full description of the landmarks in Xia et al (2020)). Descriptive statistics of these traits are given in Table 1.

**Table 1.** Descriptive statistics for tibia length and wing morphology traits measured in *Nasonia vitripennis*. For each trait, the table provides the units of measurement, the mean, the standard deviation (SD), the coefficient of phenotypic variation (CV), the minimum (Min) and the maximum value (Max).

| Traits (units) | mean |  |  | SD |  |  | CV (%) |  |  | Min. |  |  | Max. |  |  |
| --- | --- | --- | --- | --- | --- | --- | --- | --- | --- | --- | --- | --- | --- | --- | --- |
|  | G169 | G172 | combined | G169 | G172 | combined | G169 | G172 | combined | G169 | G172 | combined | G169 | G172 | combined |
| Tibia length (µm) | 603 | 618 | 610 | 42.17 | 30.09 | 38.31 | 6.99 | 4.86 | 6.28 | 401 | 461 | 401 | 692 | 680 | 692 |
| Wing length (µm) | 1921 | 2009 | 1958 | 118.52 | 88.67 | 115.42 | 6.17 | 4.41 | 5.89 | 1392 | 1516 | 1392 | 2180 | 2181 | 2181 |
| Wing width (µm) | 874 | 898 | 884 | 57.65 | 43.98 | 53.64 | 6.59 | 4.90 | 6.07 | 644 | 658 | 644 | 999 | 981 | 999 |
| Second moment area (mm<sup>4</sup>) | 0.43 | 0.51 | 0.46 | 0.10 | 0.09 | 0.10 | 23.42 | 16.98 | 21.92 | 0.12 | 0.16 | 0.12 | 0.70 | 0.86 | 0.86 |
| Aspect ratio (-) | 2.20 | 2.24 | 2.21 | 0.04 | 0.03 | 0.04 | 1.60 | 1.56 | 1.80 | 2.09 | 2.12 | 2.09 | 2.36 | 2.35 | 2.36 |

### DNA Isolation and sequencing

After detaching the wing and tibia, the remaining body of each individual female was immediately put into an Eppendorf tube with DNA buffer provided by QIAamp DNeasy® 96 Blood & Tissue Kit (Qiagen, Venlo, The Netherlands) to prevent DNA degradation. DNA was extracted by using QIAamp DNeasy® 96 Blood & Tissue Kit, following the manufacturer’s instructions. Afterwards, DNA yield and quality were checked by full-spectrum spectrophotometer NanoDrop 2000 (Thermo Scientific, Waltham, MA, USA) and Qubit 2.0 fluorometer (Invitrogen, Carlsbad, CA, USA). After qualification and quantification, DNA samples were used for Genotyping-by-Sequencing (GBS) to identify single nucleotide polymorphisms (SNPs) across the genome. GBS is a reduced representation approach that uses restriction enzymes to fragment the genome (Elshire et al., 2011), followed by size-selection and sequencing. Here, each individual DNA samples was digested by *ApeKI* restriction enzyme and adapters were ligated to both ends of the DNA fragments (one end containing a barcode to identify the individual sample, the other without). After samples were pooled and cleaned, fragments with different adapters at both ends, and with a size between 170bp and 350bp were amplified by polymerase chain reaction (PCR); and sequenced on the Illumina HiSeq X Ten platform (150 bp paired-end, Illumina, San Diego, CA, USA).

### Alignment, variants calling and filtering

After reads were sorted by individual and DNA barcodes were removed, cleaned sequence reads were aligned to the *N. vitripennis* reference genome (Nvit_2.1, https://www.ncbi.nlm.nih.gov/assembly/GCF_000002325.3/) using the BWAmem algorithm (Version 0.7.15; Li & Durbin, 2009). The alignment files were converted to BAM format, sorted, and indexed using Samtools (Version 0.1.19; Li et al., 2009). Freebayes (Version 1.9; Garrison & Marth, 2012) was used to detect genotyping variants. In total, 8,405,551 polymorphisms were identified, but the vast majority of mapped reads were spurious resulting from the inherently imperfect size selection step in creating the reduced representation library for GBS (Gileta et al., 2020). The result of this imperfect size selection is that off-target positions in the genome can be included, which we mitigated by strict selection for on-target positions, and filtering based on high genotyping rate per SNP. Filtering was done using PLINK (Version 1.9; (Purcell et al., 2007) according to the following criteria: 1) read depth between 9 and 300, 2) call rate greater than 80%, 3) Hardy-Weinberg equilibrium (HWE) exact test p-value above 10^-4^, and 4) minor allele frequency (MAF) higher than 0.02. Moreover, individuals with more than 30% missing SNPs were removed. After filtering, 8,639 SNPs remained across the whole genome of 1,230 individuals (717 in G169 and 513 in G172).

### Estimation of GEBVs with GBLUP model

Genomic best linear unbiased prediction (GBLUP) was applied to predict GEBVs. GBLUP estimates the breeding values assuming that each marker explains an equal proportion of the total genetic variance (Meuwissen 2001; VanRaden 2008). Thus, the statistical model used for GBLUP was as follows:

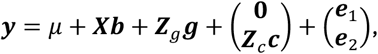

where ***y*** is the vector of phenotypic records, *μ* is an intercept, ***b*** is a vector of fixed effects, ***X*** is a design matrix relating observations to the corresponding fixed effects. ***Z***_***g***_ is an incidence matrix that relates additive polygenic values (“breeding values”) to the animals, ***g*** is a vector of random additive polygenic effects of all individuals. **0** is a vector of zeros for the individuals in G169, ***c*** is a vector of random host effects for the individuals in generation G172, ***Z***_***c***_ is the corresponding incidence matrix, ***e***1 is a vector of random residuals for G169, and ***e***2 is a vector of random residuals for G172.

The additive and host effects were assumed to be normally distributed, as ***g***∼*N*(0, ***G****σ_g_*^2^) and ***c***∼*N*(0, *Iσ_c_*^2^), respectively, where *σ_g_*^2^ and *σ_c_*^2^ are the additive genetic, and host variances, and ***G*** is a matrix of additive genomic relationships between individuals. Allele frequencies of the current population were used to construct ***G***, following Method I of VanRaden (2008), 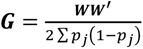, where *p_i_* is the allele frequency at locus *j*, ***W*** is a matrix of centered allele counts, with elements *W_ij_* being the code of the genotype at locus *j* for individual *i*, with (0 − 2*p_j_*) for the homozygote, (1 − 2*p_j_*) for the heterozygote, and (2 − 2*p_j_*) for the opposite homozygote. Because the host effect could not be fitted for individuals from G169, variation originating from the host may end up in the residual variance for these individuals. For this reason, we estimated the residual variance for the G169 and G172 populations separately. The residual effects of G169 and G172 were assumed to be normally distributed, 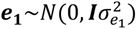 and 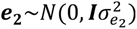, respectively, where 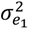 and 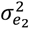 are the residual variances of G169 and G172, respectively.

Because of the large host effect in *N. vitripennis*, especially for wing and body size traits, it is essential to include host effect when estimating genetic parameters for size-related traits (Xia et al., 2020). Although we included data from both generations in the GBLUP, we estimated the genetic parameters for G172 only for which we could also estimate host effects. Heritabilities were calculated as:

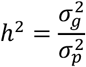

where 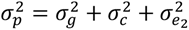 is the phenotypic variance of G172.

All analyses were performed using ASReml 4.0 (Gilmour et al., 2012), and ***G*** matrix was calculated using Calc_grm (Calus & Vandenplas, 2016).

### Population structure and linkage disequilibrium estimation

The accuracy of GEBV predictions depends on the allele frequencies and the amount of recombination between loci in the sampled population. We therefore determined the presence of population structure across the sampled generations of the *N. vitripennis* HVRx population. We estimated the population structure among the 1,230 individuals of G169 and G172 by performing principal components analysis (PCA) based on the G matrix (Remington et al., 2001). Note that pedigree-based relationships between individuals were not available in this study. To visually inspect the presence of population structure, we visualised the genomic relationships between all individuals by plotting a heatmap of the ***G*** matrix, using superheat in R 4.4.1 (Barter & Yu, 2018; Team, 2024). Analysis of the ***G*** matrix was done using ASRgenomics 1.1.3 in R (Gezan et al., 2021).

The amount of recombination between loci was determined by estimating the extent of linkage disequilibrium (LD) between SNPs as *r*^2^, using PLINK (Version 1.9; Purcell et al., 2007). The decay of LD was estimated by fitting the equation *r*^2^ = 1⁄(1 + *px*) for every focal SNP (using the nls method in R 4.4.1(Team, 2024), where x denotes the distance to every other SNP).

### Validation

To assess the accuracy of genomic prediction, we performed cross-validation. In the cross-validation process, we didvide the data set into two groups: a validation group where the phenotypes of the individuals are masked, and a training group where both phenotypes and genotypes are used for genomic prediction. The training group was used to predict the GEBVs for individuals in the validation group, and the quality of genomic prediction is measured by the correlation between the GEBVs and the observed phenotypes in the validation group.

In this study, we performed two different ways of validation: across-generation validation and 5-fold cross validation using the entire population. In the across-generation validation, the phenotypes of one generation were masked and the GEBVs of these animals were predicted using the information from the other generation. Two different scenarios were applied here: 1) across generation forward validation (AGFV), using G169 to predict G172; 2) across generation backward validation (AGBV), using G172 to predict G169. For the 5-fold cross-validation strategy, we randomly divided the population into five groups. In each cross-validation round, one group was taken as validation group and the remaining four groups as training population. The phenotypes in the validation group were masked and the training group was used to predict GEBVs of individuals in the validation group. After five rounds, all individuals in all groups received predicted GEBVs. This procedure was repeated 50 times for each trait separately, and the results were averaged over 50 random divisions of the population into a validation and a training group.

The expected value of the correlation between the GEBV and the phenotype of the validation individuals is equal to the product of heritability and the accuracy of the GEBV. Therefore, we estimated the accuracy of the GEBV by dividing the correlation between GEBVs and the observed phenotypes of individuals in the validation population by the square root of the heritability of the trait. In addition to the accuracy of genomic prediction, we also evaluated the possible dispersion in the prediction of GEBVs. The dispersion is measured by the regression coefficient of phenotypic observation on predicted GEBVs in the validation population. A value of 1 is theoretically expected for correct estimates of GEBV (no dispersion). A deviation from 1 indicates dispersion: values greater than 1 indicate underdispersion of GEBVs, whereas values smaller than 1 indicate overdispersion.

## Results

### Population structure and linkage disquilibrium (LD)

Population structure across G169 and G172 was assessed by PCA based on filtered SNPs. The PCA shows that the individuals from the two generations overlap almost completely, and the first two PCs explained only a small proportion of the total genetic variance, 6.13% and 5.27% respectively (Figure 1A). This suggests that individuals of generations G169 and G172 had very similar genotypes. The genomic relationships between all individuals were visualized using a heatmap of the G matrix (Figure 1B). The heatmap suggests that most individuals were relatively unrelated, although some individuals had a higher relationship with each other. The average genomic relationship coefficient is close to zero: 0.000783 (s.d=.113) but has a wide range (min: -0.474, max: 1.49, Supplementary Figure 1). This result is not unexpected because some individuals were collected from the same host and are therefore full or half-sibs with a high identity-by-descent. To investigate the linkage disequilibrium (*r*^2^) in our population, we plotted *r*^2^ as a function of physical distance (Figure 2). We observed a slow decay of LD, with a half-decay distance of 34.1 kb, and an average pairwise *r*^2^value of 0.47 (s.e.= 0.0017) between loci separated up to 1000 Kb.

**Figure 1.**
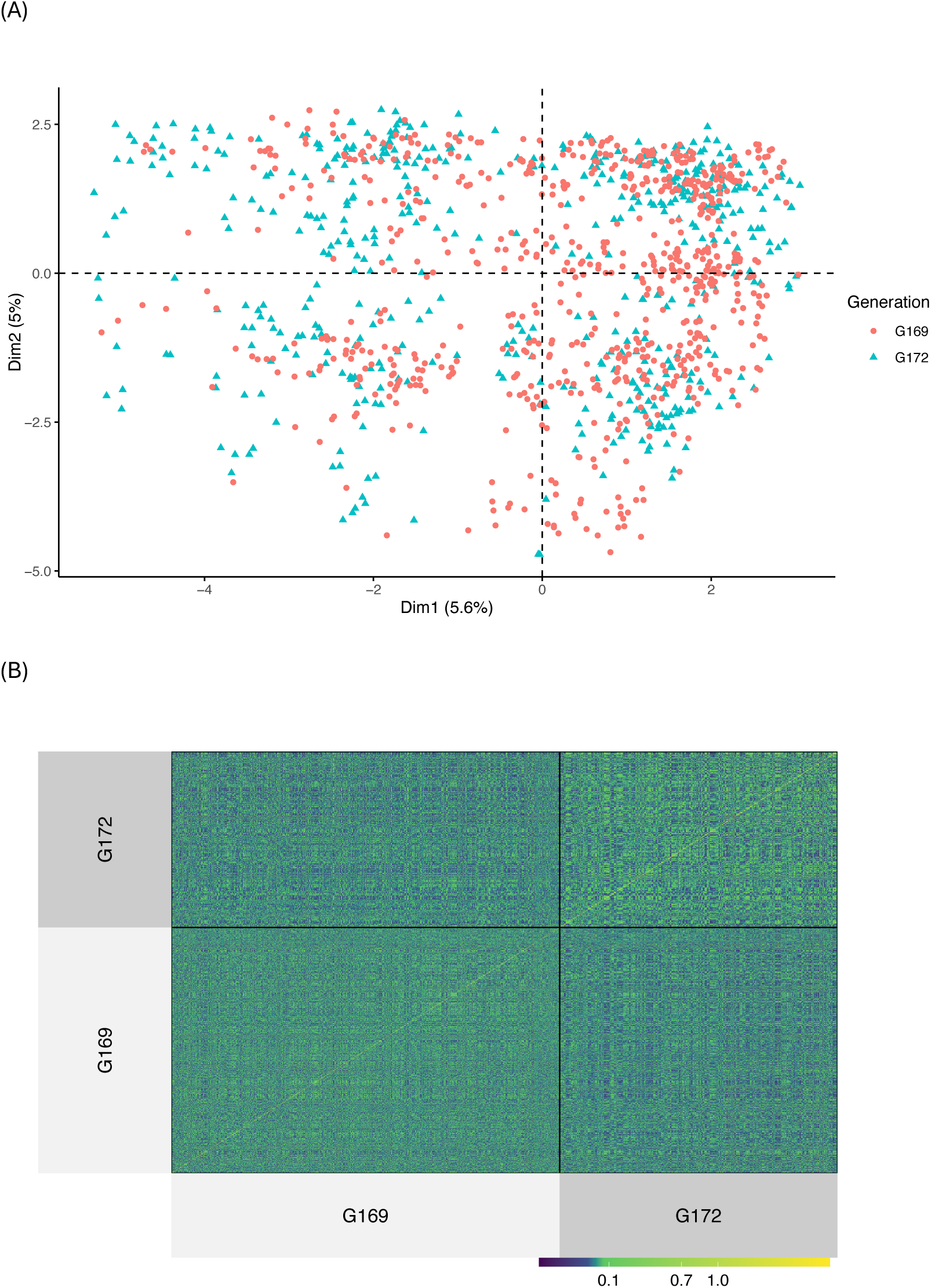
Genetic structure across G169 and G172 and relatedness among all individuals: (A) Principal component analysis (PCA) of G169 and G172, the first two principal components explained 5.6% and 5.0% of variation, respectively. Red dots represent individuals in G169, blue triangles represent individuals in G172. (B) The heatmap of the ***G*** matrix, in which high values (green) indicate a close genomic relationship between two individuals, and values of zero (dark blue) indicates no genomic relationship.

**Figure 2.**
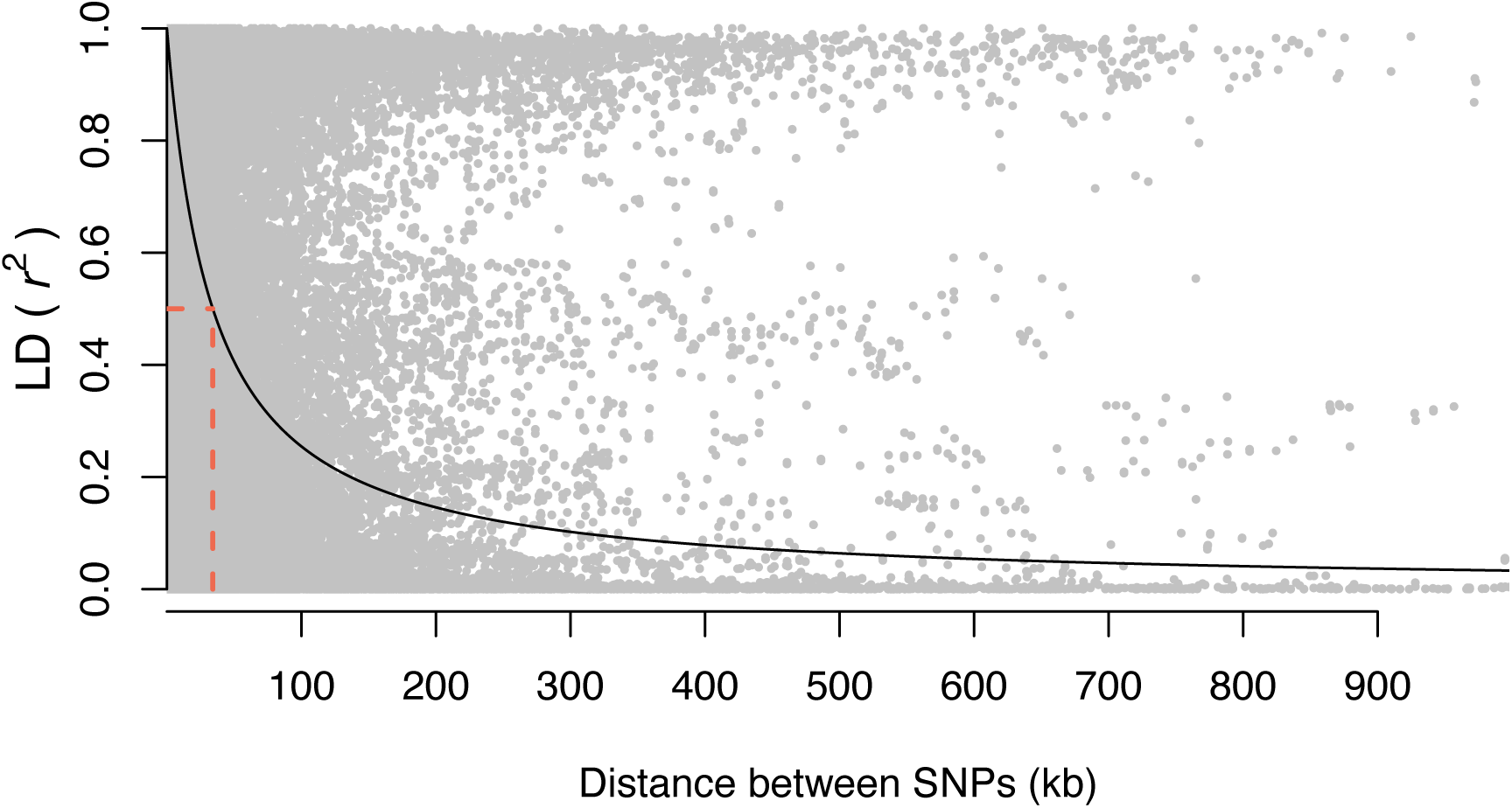
Decay of linkage disequilbrium (LD) with physical distance. Points represent the *r*^2^for pairs of markers, solid black line gives the non-linear least squares fit of *r*^2^on the distance between pairs of SNP. Dashed line indicates the half-decay LD distance at 34.1 kb.

### Validation of GEBVs

The heritability of all examined traits was moderate, ranging from ℎ^2^ =0.20 for wing width, to ℎ^2^=0.27 for second moment area (Table 2). For size-related traits, which are tibia length, wing length, wing width and second moment area, the 5-fold cross validation on the full data had a considerably higher accuracy than the across generation cross validations, either forward or backward in time. For both across generation forward and backward validation, the accuracies of genomic prediction were close to zero for the size related traits, indicating little predictive power. Remarkably, all estimated accuracies for size-related traits in across generation forward validation were negative, whereas being positive in across generation backward validation. The dimensionless wing aspect ratio showed much higher accuracy for both across generation validations, 0.47 in across generation forward validation and 0.65 in across generation backward validation. The GEBV for all traits showed higher accuries across all traits in the 5- fold cross-validation, with an average of 0.59 (s.e.=0.03).

**Table 2.** Accuracy and dispersion^1^ of genomic prediction for tibia length and wing morphology traits in *Nasonia vitripennis*, using three ways of validation with GBLUP model: across generation forward validation, across generation backward validation, and 5-fold cross-validation. Heritability (ℎ^2^) is estimated for generation G172, based on the complete data set of both generations.

| Traits | $h^2$ | across generation forward | | across generation backward | | 5-fold cross-validation | |
| --- | --- | --- | --- | --- | --- | --- | --- |
|  |  | validation |  | validation |  |  |  |
|  |  | Accuracy | Dispersion | Accuracy | Dispersion | Accuracy | Dispersion |
| Tibia length | 0.21 | -0.22 | -0.21 | 0.21 | 2.21 | 0.52 | 0.96 |
| Wing length | 0.22 | -0.12 | -0.12 | 0.17 | 1.72 | 0.60 | 1.18 |
| Wing width | 0.20 | -0.06 | -0.06 | 0.22 | 1.62 | 0.67 | 1.19 |
| Second moment area | 0.27 | -0.11 | -0.11 | 0.06 | 0.34 | 0.61 | 1.07 |
| Aspect ratio | 0.22 | 0.47 | 0.73 | 0.65 | 1.48 | 0.55 | 0.79 |

<sup>1</sup> A value of 1.0 of the regression coefficient represents the correct dispersion.

The regression coefficients of phenotypic observations on the GEBVs indicate possible dispersion in the prediction of the GEBVs. The regression coefficents of the across generation validations (forward and backward) deviated considerably from one (Table 2). However, because the accuracies of the GEBVs in both validations were close to zero, this does not provide meaningful information. In contrast, the regression coefficients for all traits were close to one in the 5-fold cross-validation (Table 2), which indicates correct estimation of GEBVs and little dispersion.

## Discussion

The need for genetic improvement of biocontrol agents has been pointed out by several authors. Despite its great success in livestock and plant breeding, genomic prediction has only been sparsely used in insects (Bernstein et al., 2023). Here we performed genomic prediction for wing morphology traits using a GBLUP model in *N. vitripennis*, on a total of 1,230 individuals with GBS-based genotypes for 8,639 SNPs. Promising accuracies were observed for the GEBVs of all traits in the 5-fold cross validation, but not both across generation validations (forward and backward). The regression coefficients in the 5-fold cross-validation indicated correct estimation of the GEBVs for all traits. Both accuracies and dispersion were in the range of those reported for honeybees (Bernstein et al., 2023).

The large difference between accuracies using different validation strategies and between traits seems to be related to the magnitude of the host effect. For size-related traits, host effects explained between 34- 56% of the phenotypic variance, whereas for wing aspect ratio they explained only 8% (Xia et al., 2020). We did not observe large host effect for wing aspect ratio because it was defined as the ratio of wing length and width, and thus the host effects on them had been scaled out. Moreover, including the host effect in the estimation of the genetic parameters (see Material and Methods) appeared insufficient to uncover the standing genetic variation of the traits that are masked by strong variation in the host environment (Xia et al., 2020).

In AGFV, we had host information only for G172. Thus, host effects were absent in the training population. In AGFV, negative accuracies were observed which suggests a false prediction signal. As the accuracies are Pearson correlation coefficients of GEBVs and the observed phenotypes of individuals in the validation population, their values can range from −1 to 1. Thus it is possible to observe negative accuracies. Nevertheless, the minimum accuracy of genomic prediction is expected to be close to zero, occurring for example when a trait is not heritable or the training population is small. Therefore, the negative accuracies found with across generation forward validation suggest a flaw in the model, likely related to the missing host effects.

In across generation backward validation, the host information of G172 was included in the training population. It was therefore surprising to observe such low accuracies for size traits. Previous studies have reported that the accuracy of genomic prediction is determined by several factors, including the number of individuals in the training set population, the heritability of the trait, and the relatedness between training and validation group (Daetwyler et al., 2008). However, we did not observe a low accuracy for all traits in across generation backward validation, with wing aspect ratio having an accuracy of 0.65. This suggests that the number of individuals in the training population and the relatedness between training and validation group are not causing observed low accuracies in the other traits. We therefore also checked the heritabilities for size traits in the G172 generation separately. When only G172 was included in the analysis, surprisingly low heritabilities for size traits were observed (Supplementary Table S1). These heritabilities are much lower than the values estimated from the combined data set, ranging from ℎ^2^ =0.20 for wing width, to ℎ^2^=0.27 for second moment area (Table 2). Furthermore, although we observed a lower heritability of wing aspect ratio in the separate G172 data set, it was not as much as for the size traits. Therefore, we conclude that the low heritability for size traits is likely the main cause for their low accuracies.

In contrast, in the 5-fold cross-validation we observed relatively high accuracies for all traits, ranging from 0.54 to 0.68. This finding further supports our previous conclusion that it is important to include host effects in the model for size-related traits (Xia et al., 2020), since in the 5-fold cross-validation we included host effects for G172 in the prediction model and estimated heterogeneous residual variances for the two generations. The accuracies in the 5-fold cross-validation are in the same range as those found in 5-fold cross-validation for several traits in the honeybee (0.44 to 0.65 for beekeeper workability traits, such as gentleness, calmness and swarming drive (Bernstein et al., 2023). VanRaden et al. (2009) compiled the accuracy of genomic prediction for a number of traits in dairy cattle, and found an average accuracy of about 0.7.

Compared to these genomic prediction studies in honeybees and dairy cattle, the accuracies in our study are very promising, given that we had a much smaller reference population size (less than 1000 individuals, compared to 2970 in honeybees) and only moderate heritabilities (∼0.22). The most likely explanations for the relative high accuracy are: 1) the small genome size of *Nasonia*, with only five chromosomes and a genetic size of only 446.9 cM (Niehuis et al., 2010), and 2) the high level of linkage equilibrium in our population. Together, both factors result in a limited effective number of independent chromosome segments (*M_e_*; Goddard, 2009). Daetwyler et al. (2008, 2010) derived the relationship between *M_e_* and the accuracy of genomic prediction, 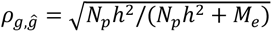, where *N_p_* is the number of individuals in the training population and ℎ^2^ is the heritability of the trait. We calculated *M_e_* following (Goddard, 2009), *M_e_* = 1⁄*Var*(*G_i≠j_*), where *Var*(*G_i≠j_*) denotes the variance among the off-diagonal elements of the genomic relationship matrix, resulting in *M_e_*= 80. Using this value in the above expression of expected accuracy, resulted in even higher accuracies (∼ 0.85) than the values found in this study. This difference may occur because the markers may not fully capture all genetic variation, and because the family structure in the population causes over-prediction of accuracy based on *M_e_* (note that the Daetwyler equation is based on a population of “unrelated” individuals). Nevertheless, the low value of *M_e_* further supports our hypothesis that the high level of LD contributes to the relative high accuracies found in this study.

In our population, LD decays relatively slowly (*r*^2^< 0.1 at 307.3 kb, and *r*^2^<0.2 at 136.5 kb). With *Nasonia* being a naturally inbreeding species, showing a high level of sib-mating (Luna & Hawkins, 2004; Grillenberger et al., 2008; Buellesbach et al., 2023), slow decay of LD is expected. However, the estimates of LD decay in our population are almost double as previous estimates based on whole genome sequencing of inbred lines from the same population (Pannebakker et al., 2020). The high level of LD and/or small *M_e_* in our population contributes to the promising accuracy in the 5-fold cross-validation because it has a relatively small effective population size (*N_e_*∼236, van de Zande et al., 2013). In applied biocontrol, the founder populations are commonly collected from the field, which may result in a large *N_e_* and thus a low level of LD and a high *M_e_*. We have not found reports on population parameters, such as *N_e_* or *M_e_*, for biocontrol agent populations so far. To maintain sufficient genetic variation, Bartlett (1993) recommended to maintain natural enemy populations with an *N_e_* >100. If the *N_e_* of a population is indeed maintained close to 100, the application of genomic prediction in biocontrol agents can benefit from a good accuracy due to the relative low *N_e_* together with the small genome size.

### Challenges in biocontrol genomic selection

In addition to the accuracy, there are some other factors that need to be taken into consideration in practical applications of genomic prediction in biocontrol agents (and insects in general). First, many biocontrol agents are insects with a small body size, which may complicate DNA extraction and genotyping of single individuals and reduce the quality of the resulting genotype data. Moreover, when DNA extraction requires the entire individual, or a large proportion of its body, it is impossible to select the parents for the next generation from the genotyped individuals. Thus, currently, genomic selection seems more suitable for insect biocontrol agents with bigger body size, so that sufficient DNA can be obtained without sacrificing the entire individual. Second, the short generation interval of many insects may limit the benefits of genomic selection in biocontrol agents. One of the main advantages of genomic selection in animal breeding is the reduced generation interval, which increases genetic gain (Meuwissen et al., 2001). However, biocontrol agents such as *N. vitripennis* already have a short generation interval, and the use of genomic selection may not allow for an even shorter generation interval. Third, small biocontrol agents usually have a short lifespan. The time required for DNA isolation, genotyping and breeding value estimation is often longer than the lifespan of the selection candidate. Thus, the application of genomic selection may be limited in organisms with a short lifespan. Yet, one could make clever use of aspects of insect biology, such as for instance the temperature dependency of lifespan on temperature which enables buying time for genotyping and parent selection (cf. Zwaan et al 1995).

A more general issue in the genetic improvement of biocontrol agents, also challenging the application of genomic selection, is the difficulty in deciding which traits to optimize (Lommen et al., 2017; Bielza et al., 2020). Kruitwagen et al. (2018) listed several candidate biocontrol traits, including high killing efficiency, robustness under (a)biotic conditions in the area of release, environmental safety, and ability to be cost-effectively (mass) reared in the laboratory. However, efficient large-scale phenotyping methods for biocontrol traits still need to be developed. Moreover, studies on the genetic basis of these biocontrol traits (i.e., heritability) also are generally lacking (Lirakis & Magalhaes, 2019). Furthermore, many biocontrol agents are parasitoid wasps, which develop their offspring in a host with a set quality. Important biocontrol traits in parasitoids, such as sex ratio, female fecundity and development time,are largely affected by the host quality (Godfray, 1994). This further emphasizes the need to account for host quality in the application of these biocontrol agents.

Finally, the genotyping methods play an important role in the application of genomic selection in biocontrol agents. Here, we applied GBS as an economical method to to obtain genome-wide marker genotypes from sequence data. A downside of GBS is the potential for relatively high rates of missing data (Poland & Rife, 2012; Crossa et al., 2013), because it produces a reduced representation of the genome, which depends on the choice of restriction enzyme (de Ronne et al., 2023). We chose the *ApeKI* enzyme after *in silico* analysis (data not shown), still our GBS protocol resulted in a lower marker density (8,639 SNPs) compared to whole-genome sequencing (205,691 SNPs; Pannebakker et al., 2020). This resulted in larger gaps between markers, potentially missing recombination events, which most likely contributed to an overestimation of our LD measurements.

An alternative to GBS and whole-genome sequencing in insects would be the development of a SNP array, which provides a cost-effective, high-throughput option for screening large SNP numbers in large sample sizes. In insects, SNP arrays have already been developed for the dengue and yellow fever mosquito *Aedes aegypti* (50K SNPs, Evans et al., 2015) and the honeybee *Apis mellifera* (>100K SNPs, Jones et al., 2020). The initial costs for desiging SNP arrays are high, but can be a robust SNP arrays can be a and economical alternative (down to €40 per sample, Suratannon et al., 2020) for large sample sizes. A promising alternative to SNP arrays in insects is using low coverage whole genome sequence data, in combination with imputation to call genotypes. When used on the Oxford Nanopore MinION platform at coverages as low as 0.1x, this method can bring the cost of genotyping down to $40 per sample and provide results within 24 h (Lamb et al., 2023).

The abovementioned factors make genomic selection for biocontrol currently a difficult and technological application. However, with the fast development of techniques, such as larger-scale phenotyping and genotyping methods, and imporved computer algorithms, the present challenges will likely be overcome in the near future. Nevertheless, our results provide a realistic assessment of the potential benefits of genomic selection when applied to natural enemies to improve biocontrol efficacy, as well as for future research. Although this study mainly focuses on application in the parasitoid *Nasonia vitripennis*, our findings and discussion of the limitations also apply to the potential application of genomic selection in insects for other uses.

## Acknowledgements

We sincerely thank Bert Dibbits, José van de Belt, Joost van den Heuvel, Jordy Litjens, Kimberley Laport and Richard Cooijmans who all contributed to the data presented in this chapter. The study is supported by a grant from the European Unions Horizon 2020 research and innovation programme under the Marie Sklodowska-Curie grant agreement No 641456.

## Author contributions

**Shuwen Xia**: conceptualization, methodology, formal analysis, investigation (lead), writing - original draft (lead). **Gabriella Bukovinszkine Kiss**: investigation, resources. **Hendrik-Jan Megens**: formal analysis. **Martien A.M. Groenen**: conceptualization; supervision, writing – review & editing. **Bas J. Zwaan**: conceptualization, funding acquisition, supervision, writing – review & editing. **Piter Bijma**: conceptualization, methodology, supervision, writing – review & editing**. Bart A. Pannebakker**: conceptualization, funding acquisition, investigation, supervision, writing - original draft, writing – review & editing.

## Supplementary Material

**Supplementary Table S1.** Estimated variance components for wing morphology traits and tibia length, based on G172 only.

| Traits | $\sigma_g^2$ | $\sigma_c^2$ | $\sigma_e^2$ | $\sigma_p^2$ | $h^2$ | $c^2$ |
| --- | --- | --- | --- | --- | --- | --- |
| tibia length ($\mu\text{m}$) | 33.15 | 644.87 | 298.29 | 976.31 | 0.03 (0.03) | 0.66 (0.04) |
| wing length ($\mu\text{m}$) | 226.12 | 6334.62 | 1987.41 | 8548.2 | 0.03 (0.03) | 0.74 (0.03) |
| wing width ($\mu\text{m}$) | 84.44 | 1597.19 | 432.22 | 2113.8 | 0.04 (0.03) | 0.76 (0.03) |
| second moment area<br>($\text{mm}^4$) | $4.60 \times 10^{-4}$ | $5.42 \times 10^{-3}$ | $1.91 \times 10^{-3}$ | $7.79 \times 10^{-3}$ | 0.06 (0.03) | 0.70 (0.04) |
| aspect ratio (-) | $1.64 \times 10^{-4}$ | $1.58 \times 10^{-4}$ | $9.17 \times 10^{-4}$ | $1.24 \times 10^{-3}$ | 0.13 (0.06) | 0.13 (0.05) |

$\sigma_g^2$ : additive genetic variance; $\sigma_c^2$ : variance of host effects; $\sigma_e^2$ : residual variance of G172; $\sigma_p^2$ : phenotypic variance of G172, calculated as $\sigma_p^2 = \sigma_g^2 + \sigma_c^2 + \sigma_e^2$ ; $h^2$ : heritability of G172, calculated as $h^2 = \frac{\sigma_g^2}{\sigma_p^2}$ , standard deviation in parentheses; $c^2$ : proportion of variance due to host effects of G172, calculated as $c^2 = \frac{\sigma_c^2}{\sigma_p^2}$ , standard deviation in parentheses.

Variance components were estimated using data from G172 only, and the following statistical model:

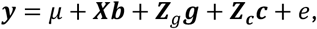

where ***y*** is the vector of phenotypic records for G172, *μ* is an intercept, b is a vector of fixed effects, ***X*** is a design matrix relating observations to the corresponding fixed effects. ***Z***_***g***_ is an incidence matrix that relates additive polygenic values (“breeding values”) to the animals, ***g*** is a vector of random additive polygenic effects of all individuals, ***c*** is a vector of random host effects for the individuals in generation G172, ***Z***_***c***_is the corresponding incidence matrix, ***e*** is a vector of random residuals for G172.

**Supplementary Figure 1.**
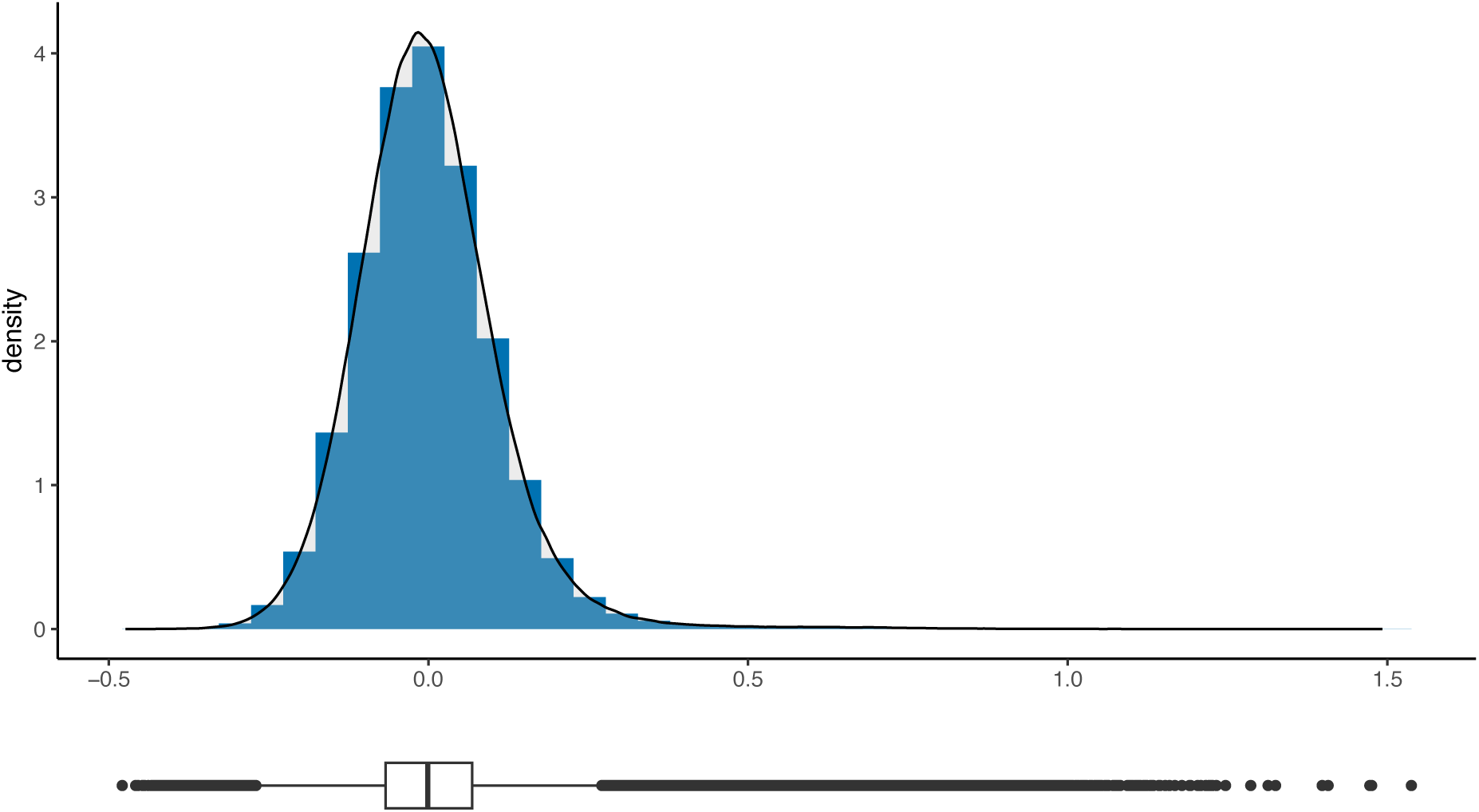
Density plot of the distribution off-diagonal values of the ***G*** matrix, indicating the genomic relationships between individuals across generations G169 and G179.

